# Number of detected proteins as the function of the sensitivity of proteomic technology in human liver cells

**DOI:** 10.1101/2021.11.24.469687

**Authors:** Nikita Vavilov, Ekaterina Ilgisonis, Andrey Lisitsa, Elena Ponomarenko, Tatiana Farafonova, Olga Tikhonova, Victor Zgoda, Alexander Archakov

## Abstract

The main goal of the Russian part of C-HPP is to detect and functionally annotate missing proteins (PE2-PE4) encoded by human chromosome 18. However, identifying such proteins in a complex biological mixture using mass spectrometry (MS)-based methods is difficult due to the insufficient sensitivity of proteomic analysis methods. In this study, we determined the proteomic technology sensitivity using a standard set of UPS1 proteins as an example. The results revealed that 100% of proteins in a mixture could only be identified at a concentration of at least 10^−9^ M. The decrease in concentration leads to protein losses associated with technology sensitivity, and no UPS1 protein is detected at a concentration of 10^−13^ M. Therefore, two-dimensional fractionation of samples was applied to improve sensitivity. The human liver tissue was examined by selected reaction monitoring and shotgun methods of MS analysis using one-dimensional and two-dimensional fractionation to identify the proteins encoded by human chromosome 18. A total of 134 proteins were identified. The overlap between proteomic and transcriptomic data in human liver tissue was ~50%. This weak convergence is due to the low sensitivity of proteomic technology compared to transcriptomic approaches. Data is available via ProteomeXchange with identifier PXD026997.

## 1. Introduction

The detection of the so-called “missing” proteins [1], the existence of which has not been confirmed by mass spectrometry (MS) or other methods, remains a major challenge a decade after the start of the Human Proteome Project (HPP).

The Russian contribution to the project is associated with the study of chromosome 18 proteins on three types of biological material: blood plasma [2], liver tissue, and the HepG2 cell line [3].

Human chromosome 18 encodes 264 genes, of which 255 have evidence at protein level (PE1) in neXtProt database. Transcripts were found for nine genes but with no confirmation at the protein level (PE2), and there is no information about whether these encode any proteins (PE5) for 11 genes. Thus, human chromosome 18 contains nine “missing” PE2 proteins, PE3 and PE4 proteins are absent [4].

The Russian group has previously studied human liver tissue to identify proteins unique to this organ and measured their concentration in normal liver and a cell line that models HepG2 liver cells. For this purpose, standard methods of targeted and shotgun MS analyses were used.

The main constraint in the study of “genome-wide proteomes” is the limitations of sensitivity and dynamic range of concentrations measured by modern proteomic technologies.

The primary reasons for the insufficient sensitivity of modern proteomic approaches are relatively low degree of ionization, which is 10–30% [5], use of nanospray ionization sources, and co-elution of numerous peptides. The latter leads to a competition for proton and the suppression and distortion of the signal of low-intensity peptide ions.

DMSO, as a chromatographic mobile phase additive, is a possible solution to the problem of low ionization because this method increases the number of identifications by 20% [6]. Complex biological sample fractionation methods may solve this problem because it can use more material and reduce the total number of peptides in one sample. Assuming that a whole cell lysate contains the complete human proteome of ~20,000 proteins, when fractionating such a sample into uniform 20 fractions, 20-fold fractionation yields a significant decrease in complexity, but we registered some minor overlapping between fractions. In this case, the enrichment of each fraction with certain peptides was also observed.

Thus, a fold increase in sensitivity was demonstrated using biochemical methods for protein and peptide pre-separation based on 1D PAGE, 2D PAGE, HPLC, and peptide isoelectric focusing (IEF) [7,8].

There are several methods of protein and peptide fractionation; however, for our task, the most appropriate method is the chromatographic separation of peptides which collects fractions during reverse-phase chromatography under alkaline conditions. Such separation is well reproduced, which allows the creation of a reliable method for selected reaction monitoring (SRM) analysis of each peptide fraction [3].

The scientific novelty of this work lies in the fact that using a standard set of UPS1 proteins it was shown that the amount of detected proteins depends on the sensitivity of the technology at a concentration of 10^−9^ M all 48 proteins are detected, and at a concentration of 10^−12^ M - only 8 proteins. When the UPS1 sample 10^−12^ M was concentrated to 10^−10^ M and MS analysis was performed, almost all proteins are detected and this allows explaining the existence of a gap (~ 50% of the proteins not detected are in biological samples in a concentration of less than 10^−10^ M). The comparison of the proteomic data and transcriptomes of three human liver tissue samples was obtained using RNA-seq Illumina HiSeq, qPCR, and ONT-Oxford Nanopore. Technology revealed the insufficient sensitivity of modern proteomic technologies. The expression of the entire human chromosome 18 transcriptome was detected [9].

In this article, it was shown that the use of two-dimensional fractionation increases the sensitivity of proteomic technologies by 2 folds in the number of identified proteins, both Shotgun analysis and SRM. Previous results of proteomic analysis of human liver samples are consistent with the results obtained in this article (60 proteins identified by SRM analysis). Two-dimensional fractionation allowed us to find 134 proteins encoded by the 18th human chromosome. However, the greatest achievement in this work is the detection in a standard set of all 48 UPS1 proteins with a sensitivity of 10^−9^ M, with a sensitivity of 10^−12^ M-8 proteins. When this mixture is concentrated with an initial protein content of 10^−12^ M by 100 times up to 10^−10^ M, it is possible to register 45 UPS1 proteins, which indicates a direct dependence of the number of proteins on the sensitivity of the technology. Under proteomic technologies, we mean not only the sensitivity of the detector of the device, but also the sample preparation for MS analysis and complexity of the composition of the biological sample.

## 2. Results

### 2.1. Shotgun (panoramic) MS analysis

The data obtained by shotgun proteomic analysis were processed using MaxQuant software with an FDR cutoff of 1%. Peptide spectrum matches (PSMs), peptides, and proteins were validated at a 1.0% false discovery rate (FDR) estimated using the decoy hit distribution; the FDRs of PSM, peptide, and protein were 0.12%, 0.63%, and 0.91%, respectively. In a non-fractionated liver sample, 2087 protein groups were identified, 25 of which were proteins encoded by human chromosome 18. Proteins that cannot be resolved were considered one group, since only common peptides were found for them. Among 2087 protein groups, only 394 could not be resolved, since they contain common peptides. The remaining 1693 proteins were identified by 2 or more unique peptides. Twenty fractions were obtained after fractionation on RP under alkaline conditions, and 4936 protein identifications were revealed by proteomic analysis, 63 of which were the proteins encoded by human chromosome 18 (Figure 1, Table S3). The analysis of the data obtained by both methods revealed the identification of 5022 proteins (Figure S1, S2), 64 of which were encoded by chromosome 18 proteins (Figure 2). Among 5022 protein groups, 387 protein groups could not be resolved, and the remaining 4635 proteins were identified by two or more unique peptides. In each liver sample, 31 proteins encoded by human chromosome 18 were detected. The differences between liver samples 1, 3, and 5 were 21, 9, and 18 protein identifications, respectively (Figure 2). Thus, the sensitivity of the shotgun method increased 2.5 times when using an additional method of peptide fractionation based on the number of protein identifications. Notably, the same trend was revealed when analyzing protein identifications for other chromosomes. The percentage coverage of each chromosome in the number of protein identifications was evenly distributed over each chromosome by ~25% (Figure S3). Thus, we can conclude that 5022 proteins in one type of cell tissue are not the limit since, in fact, a larger number of proteins are expressed. However, due to the limited sensitivity of shotgun analysis, the exact number of proteins for this type of cell cannot be determined.

**Figure 1.**
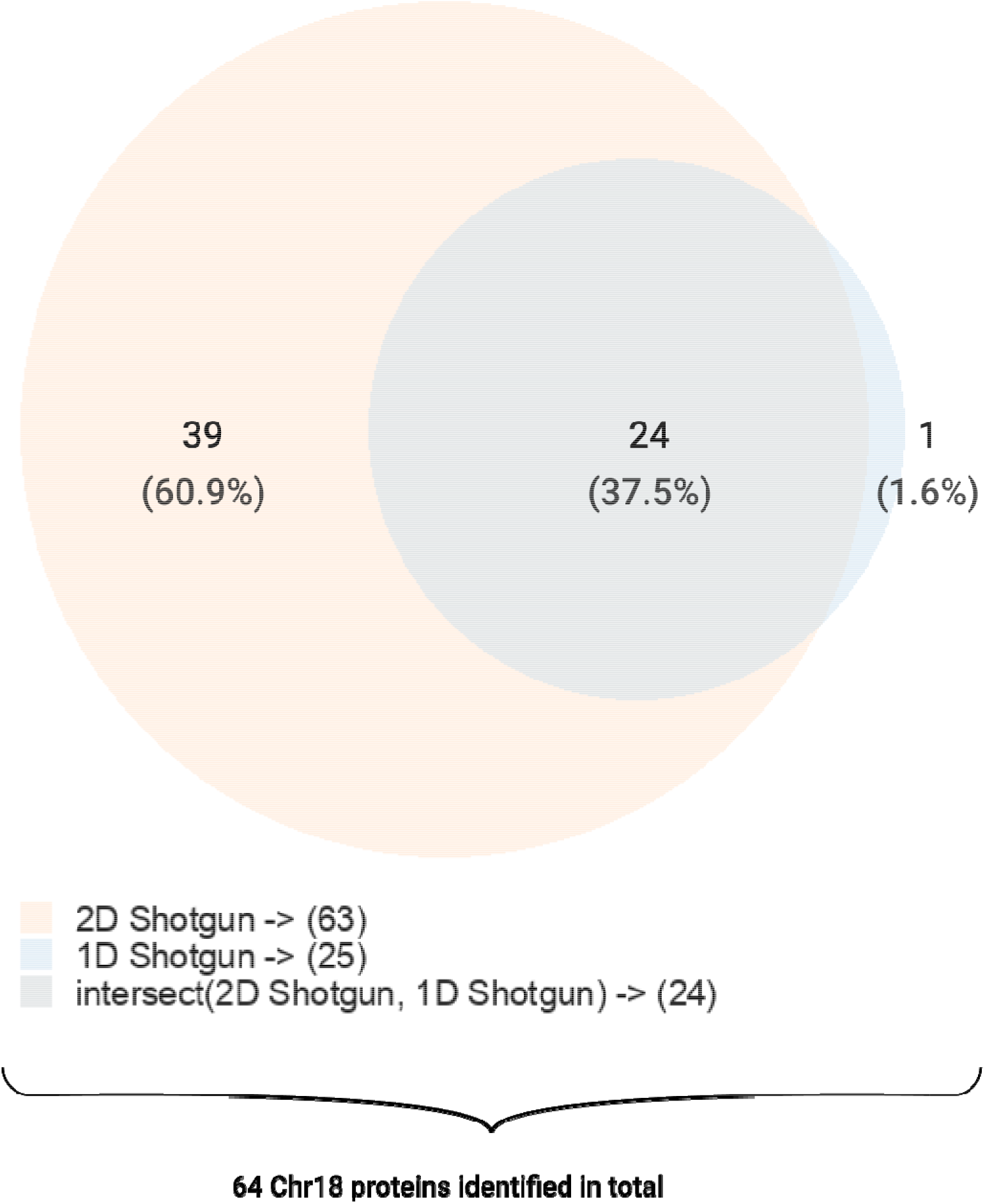
Venn diagram showing the proteins encoded by human chromosome 18 identified using one-dimensional (1D shotgun) and two-dimensional (2D shotgun) methods in human liver samples.

**Figure 2.**
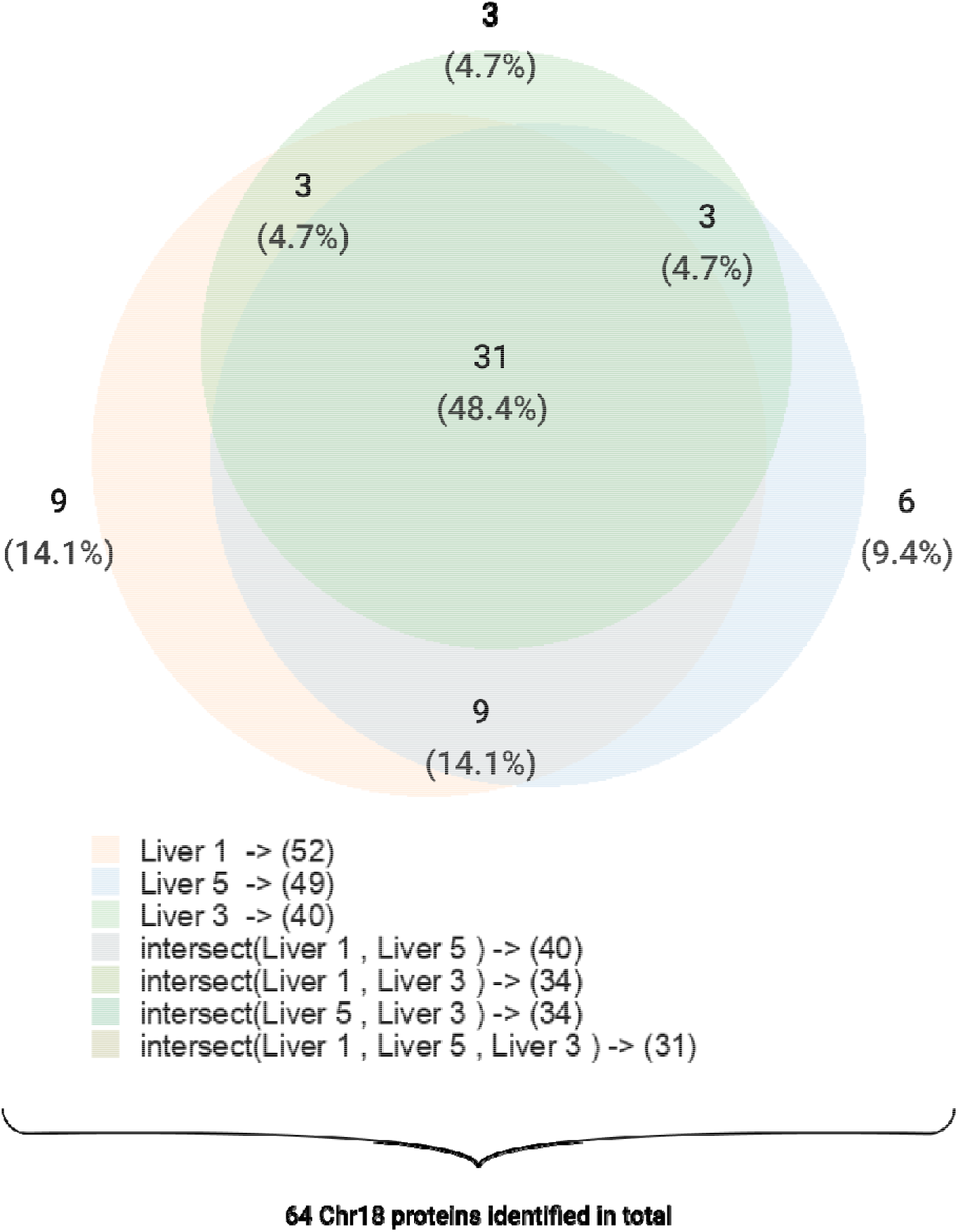
Venn diagram showing the proteins encoded by human chromosome 18 identified using one-dimensional and two-dimensional methods in each human liver sample.

### 2.2. Targeted MS analysis – SRM SIS

Targeted MS analysis is more sensitive but suffers from limited multiplexity [3]. In this experiment, 265 proteins encoded by human chromosome 18 were tested with 60 protein identifications registered in the whole sample, which was twice as high as that obtained by the shotgun (panoramic) analysis (Table S4). The analysis of peptide fractions after chromatographic separation revealed 110 proteins encoded by human chromosome 18 (Figure 3, Table S5). When using 2D technology, as a result of additional manipulations, the concentration of individual peptides may decrease and they are not detected, since the concentration becomes below the sensitivity limit. In each liver sample, 71 identical identifications were found, and the differences between samples 1, 3, and 5 were 27, 20, and 18 protein identifications, respectively (Figure 4). Targeted MS analysis remains a priority for the development of the most sensitive methods to study the undetected part of the proteome using shotgun MS analysis. An increase in sensitivity was confirmed by the minimum detectable protein concentration. In one-dimensional SRM analysis, the detection limit of a protein was 10^−12^ M, and in two-dimensional analysis, it was possible to identify proteins at a concentration of 10^−13^ M. In total, 118 proteins encoded by human chromosome 18 were found, which is 45% of the encoded by chromosome 18 proteins.

**Figure 3.**
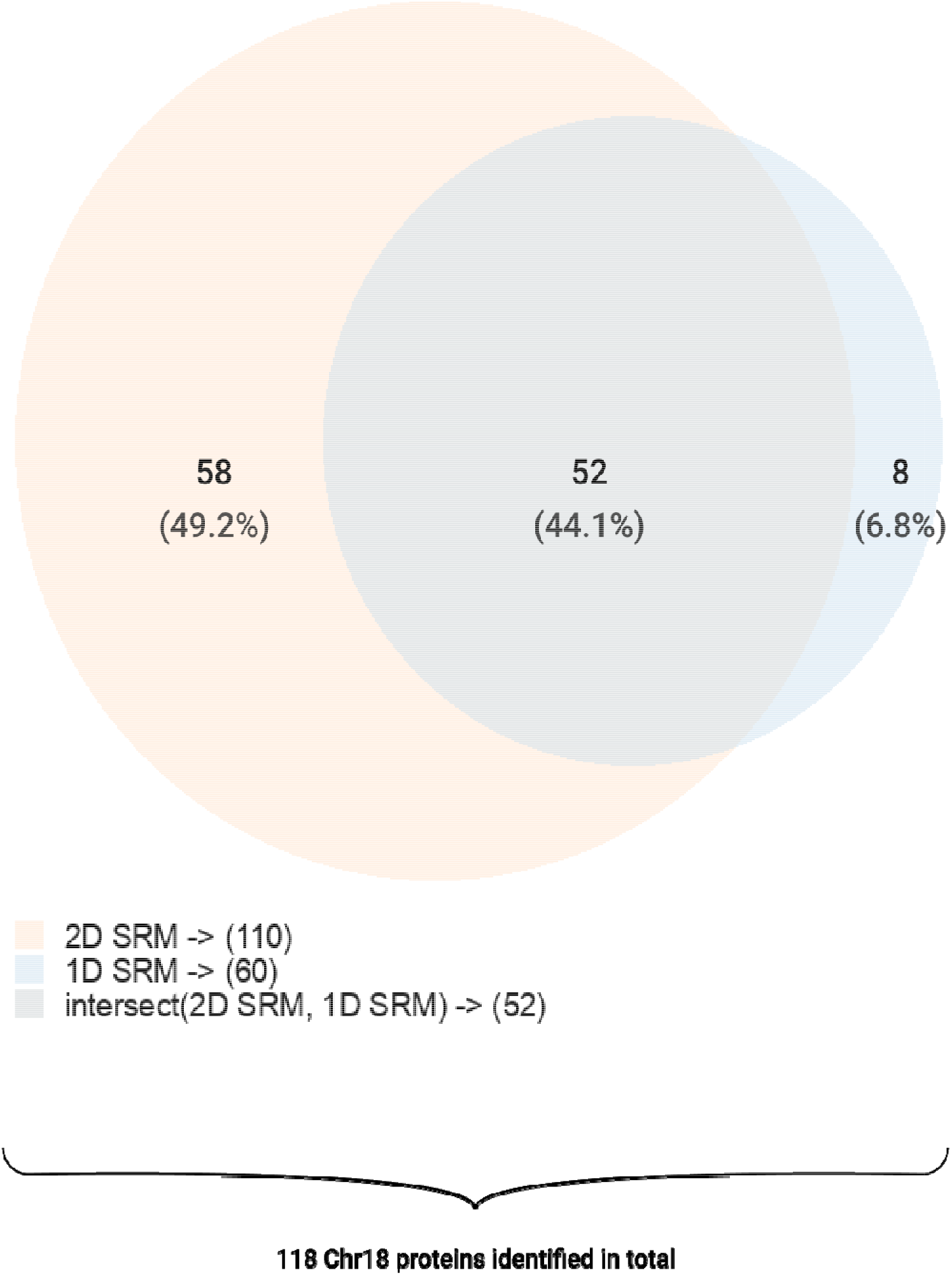
Comparison of protein identifications obtained using targeted scanning of the whole sample (1D SRM) and peptide fractions after chromatographic separation under alkaline conditions (2D SRM).

**Figure 4.**
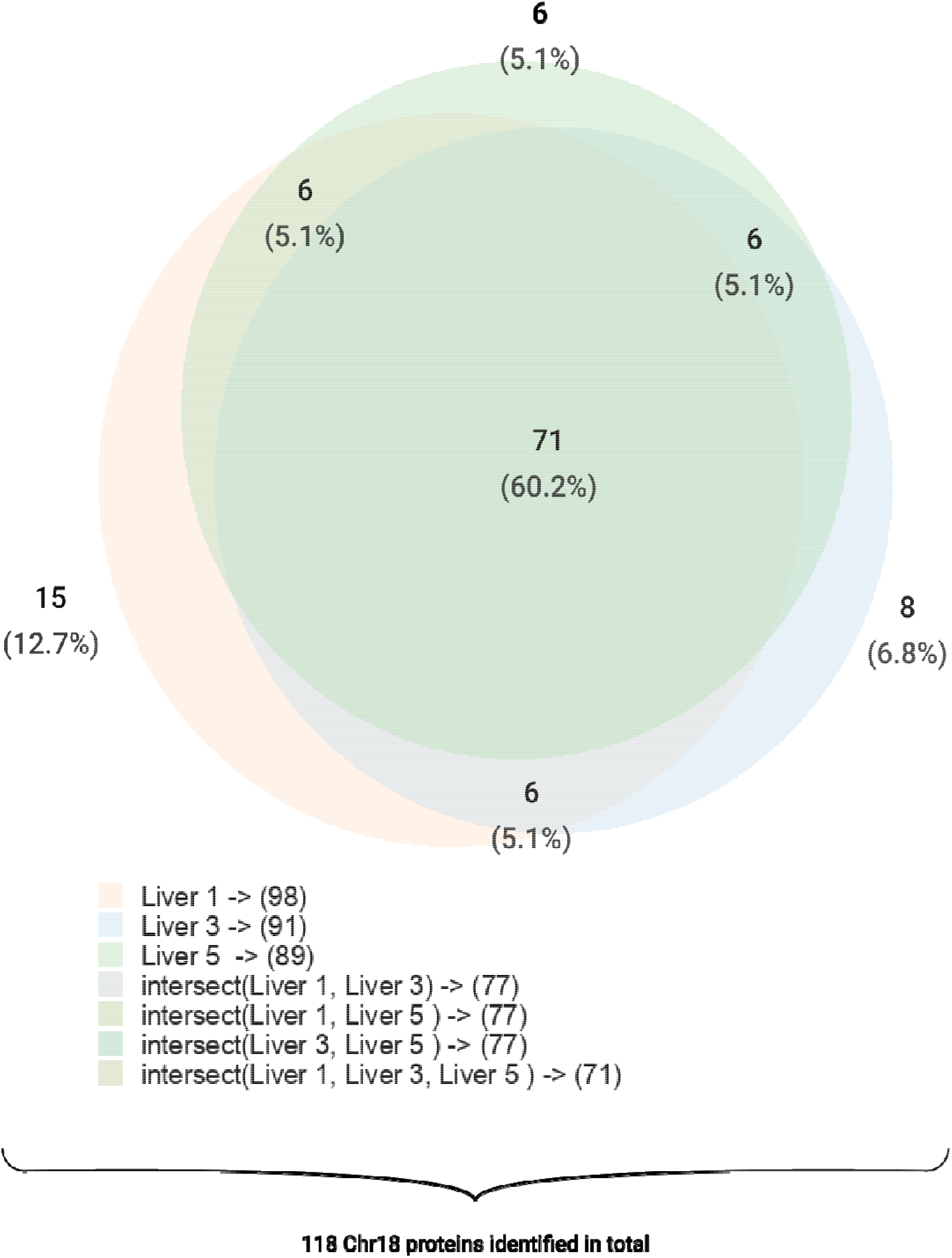
Comparison of protein identifications obtained using the analysis of each liver sample and targeted SRM method.

### 2.3. Comparison of chromosome 18 proteins identified in liver samples using the shotgun method and targeted SRM analysis

Although targeted SRM analysis is inferior to shotgun MS analysis in terms of multiplexity, it allows the registration of more chromosome 18 proteins in liver cells and measuring their concentrations. This is because during the SRM analysis on a triple quadrupole, we limit the visibility area of the device. However, when we apply the shotgun method, the device shoots at the entire possible scanning range, thereby likely increasing the dynamic range of protein concentrations registered by SRM analysis compared to the shotgun method (Figures. 5 and 6).

**Figure 5.**
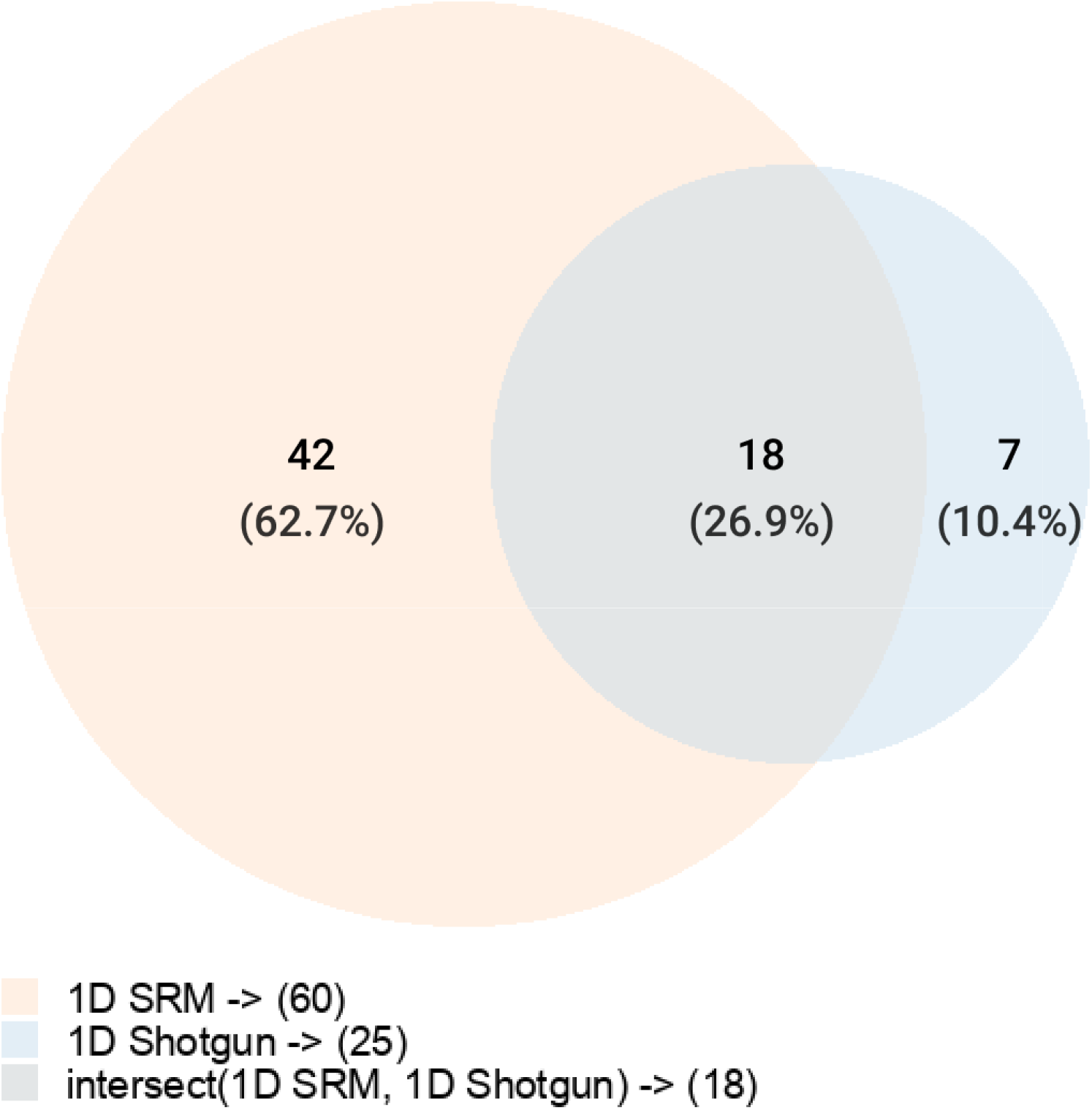
Comparison of the results of protein identification using one-dimensional shotgun and SRM methods by the number of chromosome 18 proteins identified.

**Figure 6.**
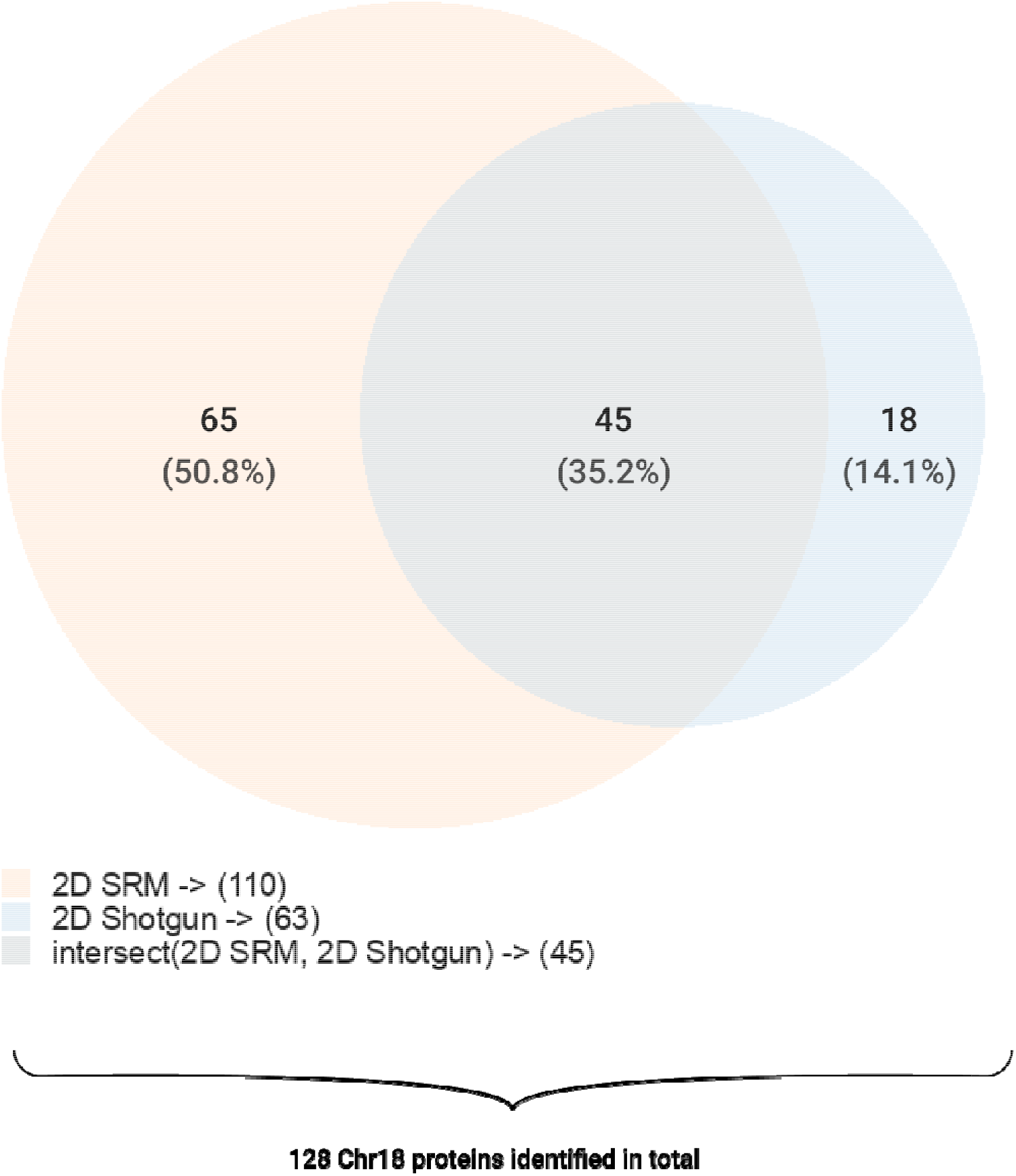
Comparison of the results of protein identification using two-dimensional shotgun and SRM methods by the number of chromosome 18 proteins identified.

Overall, we identified 134 proteins encoded by human chromosome 18. Among all the methods used, the maximum coverage of human chromosome 18 gene products was provided by two-dimensional fractionation using targeted analysis (2D SRM) with 110 proteins identified. However, the most comprehensive picture was obtained using all four methods (Figure 7). The shotgun analysis also made a significant contribution to the number of proteins identified, and 16 proteins were detected only by shotgun analysis. Thus, the peptides can be identified using shotgun analysis, and the SRM technique can be adjusted to obtain better results.

**Figure 7.**
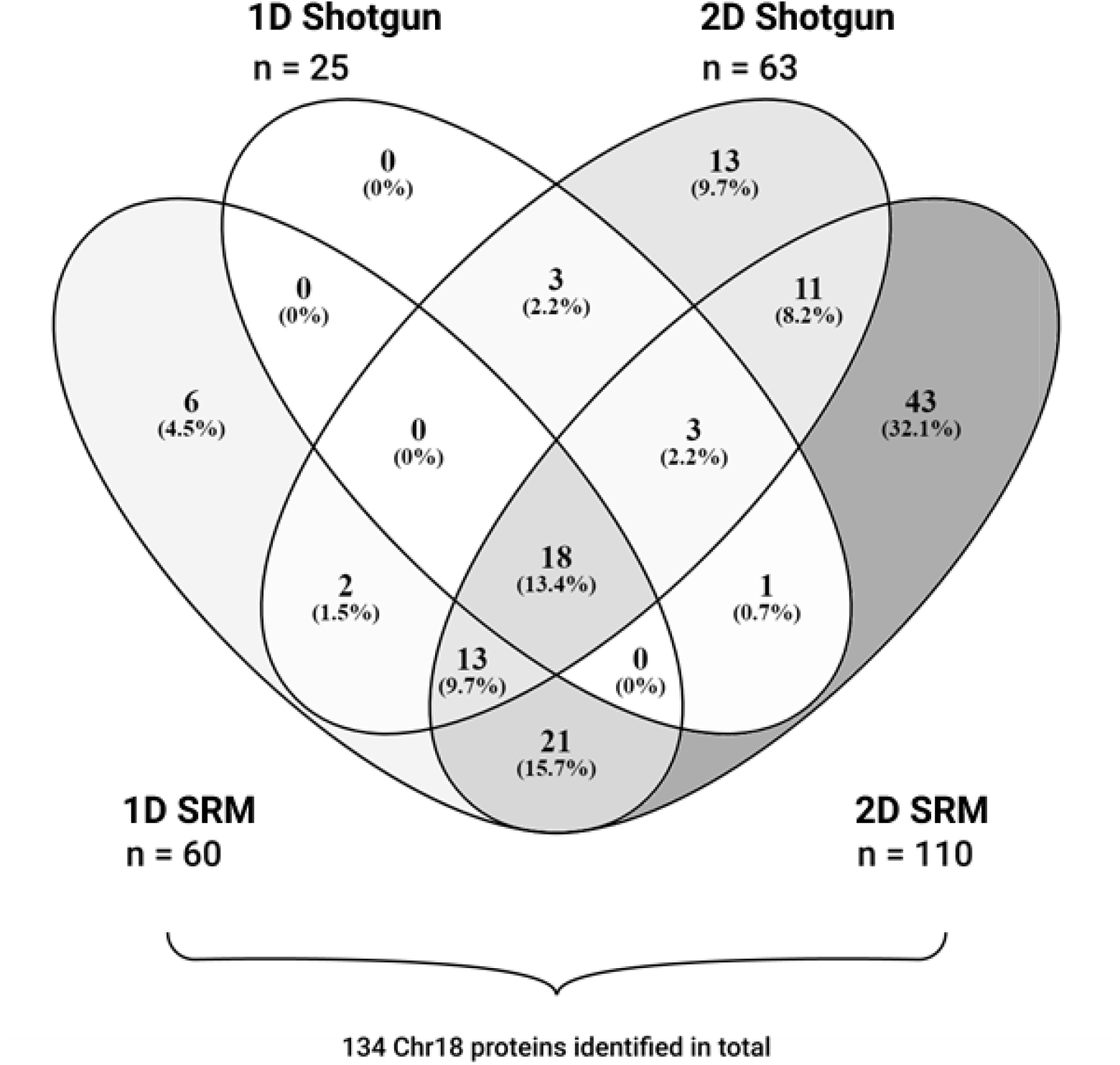
Comparison of protein identifications obtained using all methods.

The application of all scanning methods renders achieving 50% coverage of the gene products of human chromosome 18.

### 2.4 Number of UPS1 proteins detected is a function of the sensitivity of proteomic technologies

One of the reasons for the incomplete coverage of the human liver proteome is the insufficient sensitivity of MS technology when analyzing complex biological samples. To prove this, we conducted an experiment with a standard UPS1 object.

UPS1 is a set of 48 human proteins at the same concentration. The application of UPS1 was aimed at showing the part of the proteome measured by the SRM technique and comparing the results with those obtained when measuring the human proteome in human liver samples.

In the first stage, UPS1 was studied using the shotgun method to select the peptides to be used in SRM analysis, and three peptides were selected for each protein in the UPS1 set. An SRM analysis method for UPS1 set proteins was then developed based on these peptides.

UPS1 was gradually diluted until the signal disappeared in its pure solution, at a constant signal-to-noise value and a decreasing signal-to-noise ratio (an *Escherichia coli* sample was used as noise).

At UPS1 concentration of 10^−12^ M, at which the signal was sufficiently weak, a concentration of 100 times was applied for the sample with a constant signal/noise value and a pure solution. Such an approach is not possible for the sample with a decreasing signal-to-noise value, as the final concentration of *E. coli* is 2×10^−3^ M. For technical reasons, such an amount of protein cannot be used for chromatographic analysis on an HPLC column.

An experiment with a decreasing signal-to-noise value can most accurately simulate the situation with protein identification in a complex biological sample. The concentration of each protein in a biological sample is unknown; however, 100% of proteins in a sample at a concentration of 10^−9^ M can be identified using SRM analysis of the UPS1 (Figure 8).

**Figure 8.**
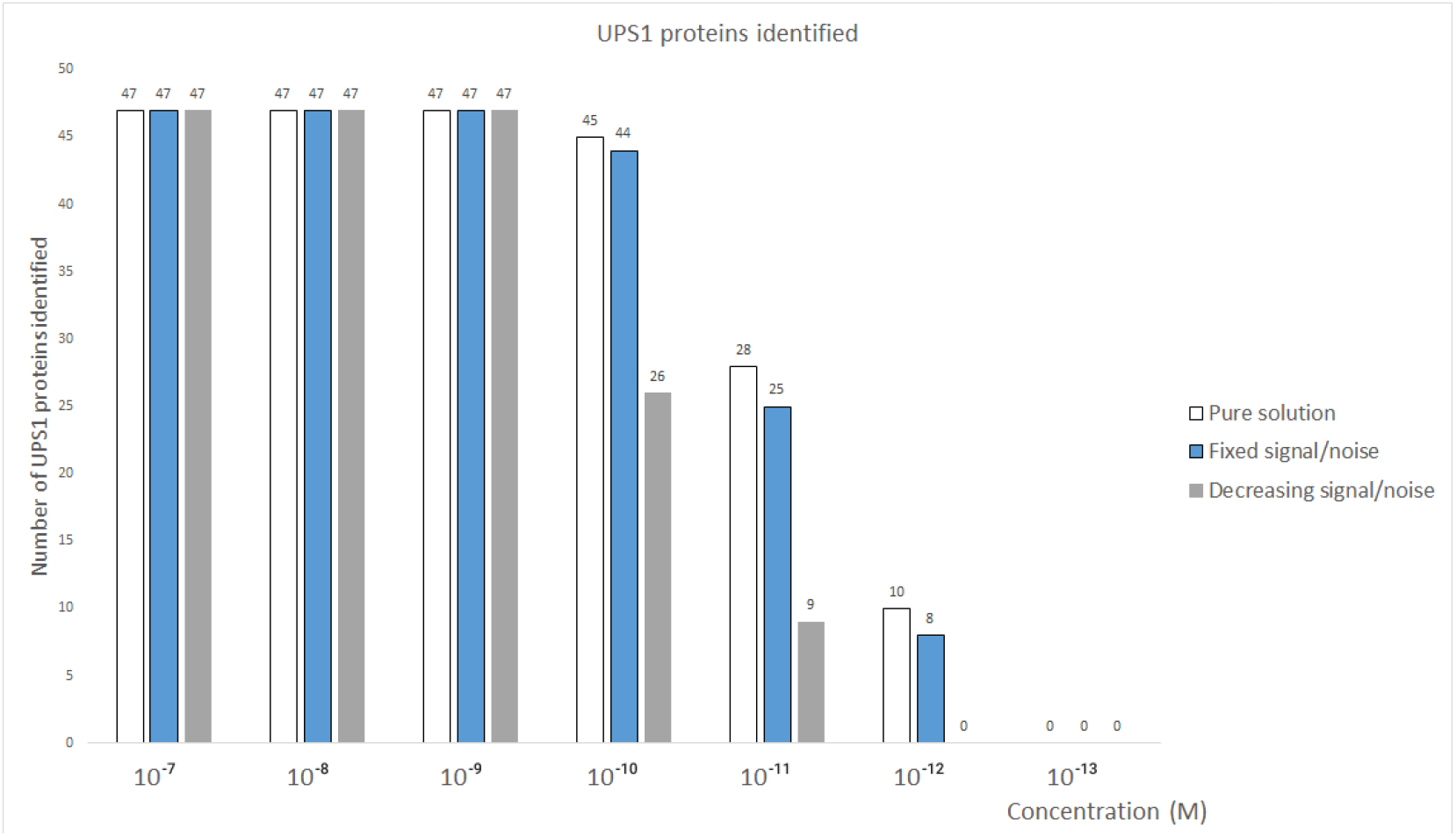
Dependence of the number of UPS1 identified on the concentration in its pure solution at a constant signal-to-noise ratio and decreasing signal-to-noise ratio.

If proteins are at a lower concentration, they might not be registered. The first losses begin when the concentration reaches 10^−10^ M, and even a single protein of the UPS1 mixture is not registered at a concentration of 10^−13^ M. When we recovered the sample that was diluted 100 times (from 10^−12^ M to a point of 10^−10^ M), we restored the number of protein identifications (Figure 9). In addition, from the experiments with sample recovery, we can conclude that adhesion to plastic and additional manipulations do not significantly contribute to the loss of protein identifications at low concentrations. The primary challenges are the limited sensitivity and dynamic range of proteomic techniques at low concentrations.

**Figure 9.**
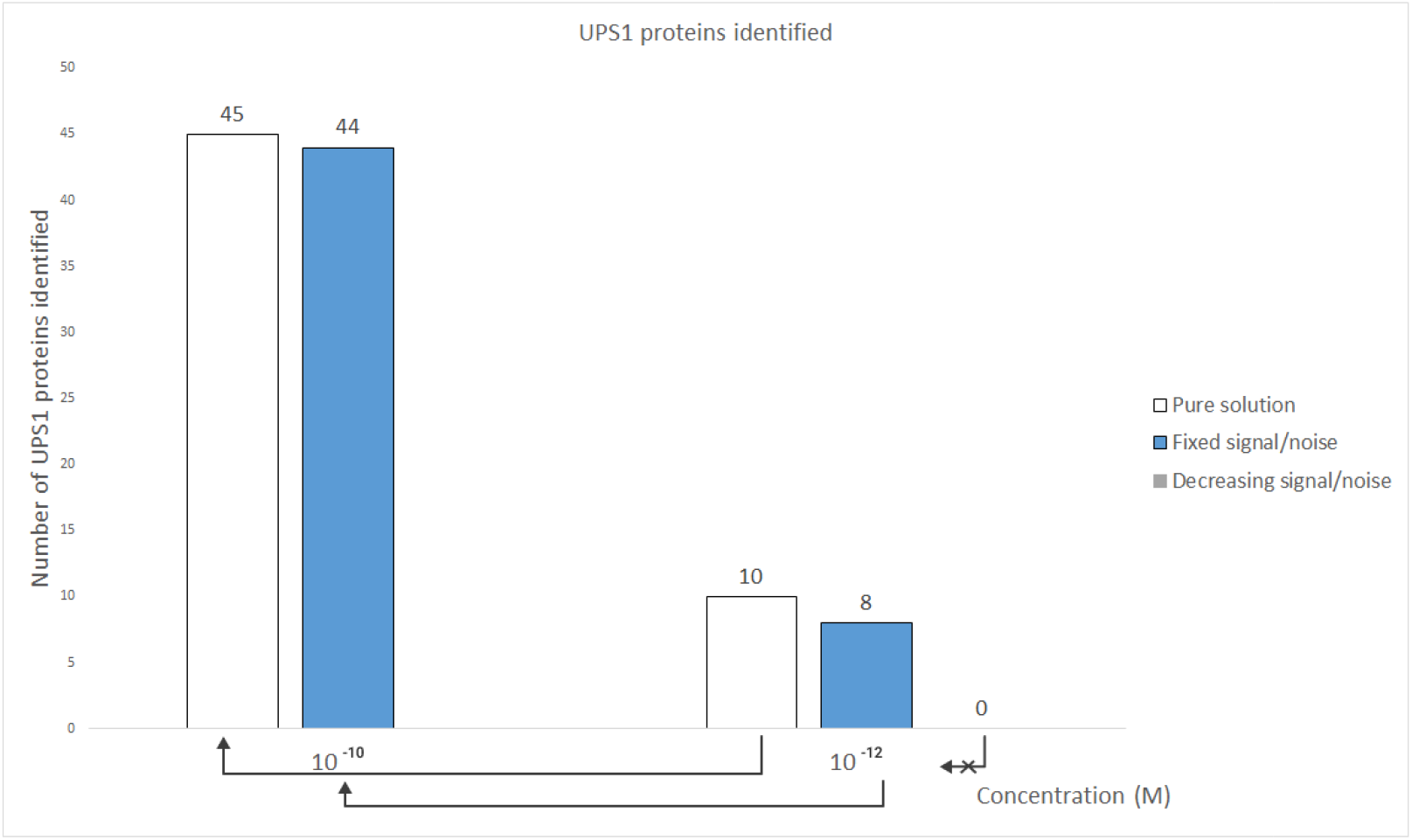
Samples UPS1 with the addition of E. coli and a pure solution at a concentration of 10^−12^ M were concentrated 100 times to 10^−10^ M and MS analysis was carried out, which showed the possibility of restoring sensitivity.

## 3. Discussion

### 3.1 Shotgun analysis

In our previous study, we investigated the proteome of the hepatocellular cancer cell line HepG2. The significant proportion of identified proteins were the same in the liver and HepG2 samples (2989, 46.3%); however, a large set of the proteins were detected only in the HepG2 cell line (1435, 22.2%) or in the liver (2033, 31.5%). According to the obtained data, the total number of proteins found in HepG2 cells and human liver tissue cells was 4424 and 5022, respectively (Figure 10). This difference could be explained by the fact that the liver tissue includes several different types of cells; therefore, some proteins registered in the liver samples may belong to them. However, the cells of the HepG2 line are a homogeneous culture, and more proteins characteristic of HepG2 were registered in these samples. However, a large set of protein identifications were common in the liver and HepG2 samples (2989, 46.3%), which shows their similarity and justifies the use of HepG2 cells as a model of human liver cells [10].

**Figure 10.**
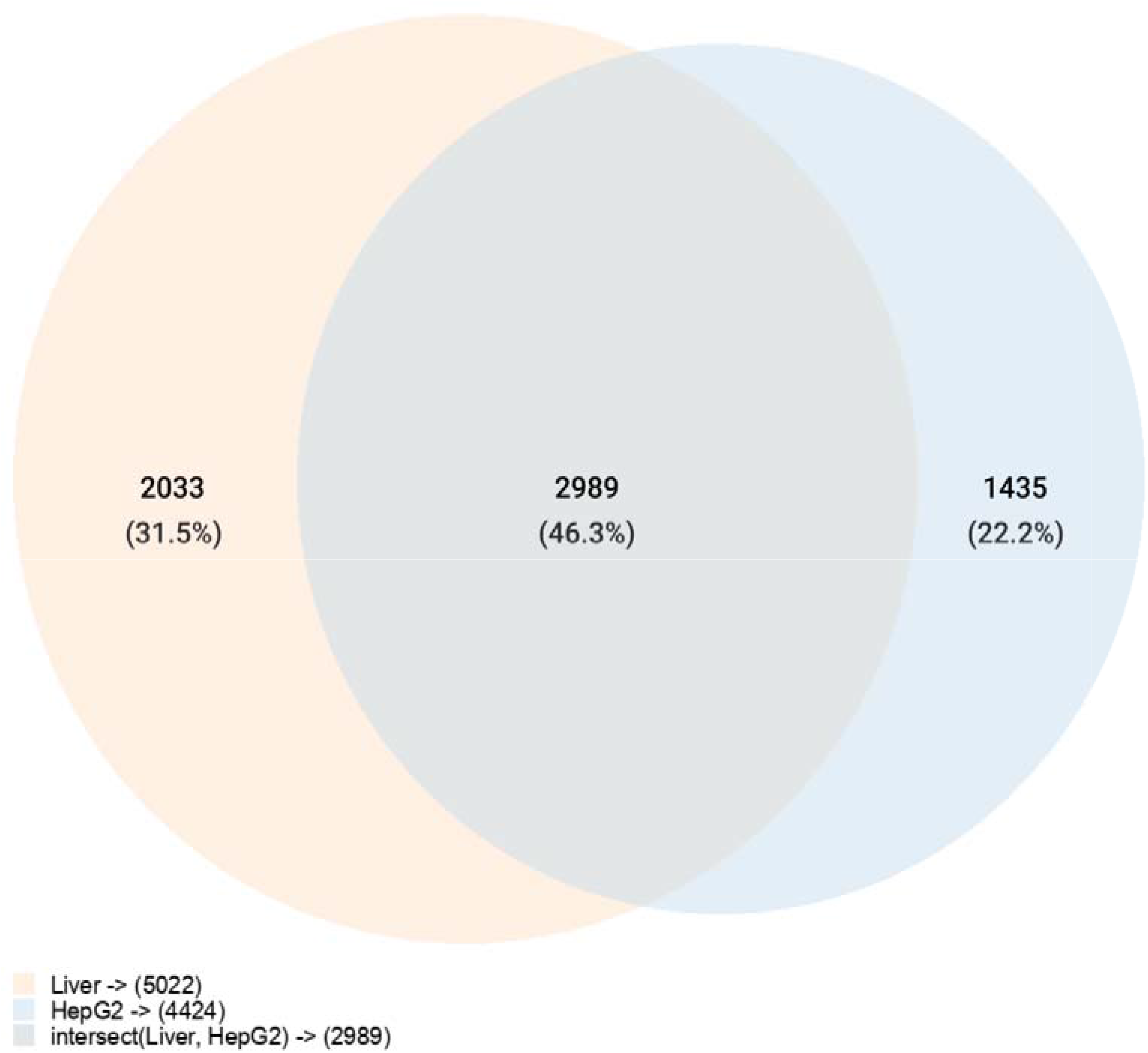
Comparison of proteins identified in human liver cells and HepG2 cells using the shotgun method.

### 3.2 SRM analysis

Previously, we studied the proteome of the hepatocellular cancer cell line HepG2 using SRM analysis. In this study, a comprehensive investigation of the proteome in the context of the dynamic concentration range provided a clearer picture that determines the similarity between these two types of samples (Figure 11). In the HepG2 cell line, only 11 (8.5%) proteins were encoded only in this type of biomaterial. This might be due to the altered metabolism of cancer cells, in contrast to normal liver cells. For example, one of these proteins is collagen and calcium-binding EGF domain-containing protein 1, which is required for embryogenesis, which may indicate a low degree of HepG2 cell differentiation compared to human liver cells [11]. Most of the proteins identified in liver cells were also identified in HepG2 (77, 59.7%). However, 41 proteins (31 8%) encoded by chromosome 18 were identified in liver cells because three biological samples were used in this analysis, and as shown above, there was an interindividual variability between these samples, which gives additional identifications in human liver samples [12]. Interindividual variability is a fluctuation around some homeostatic parameters, and in this case, it was the different concentrations of proteins. Some proteins were not identified and were present only in one human liver sample, because the concentration of these proteins was below the sensitivity limit of the technique. The device sensitivity was at the level of low fmole and amole concentrations, which correlates with data from other laboratories [13, 14].

**Figure 11.**
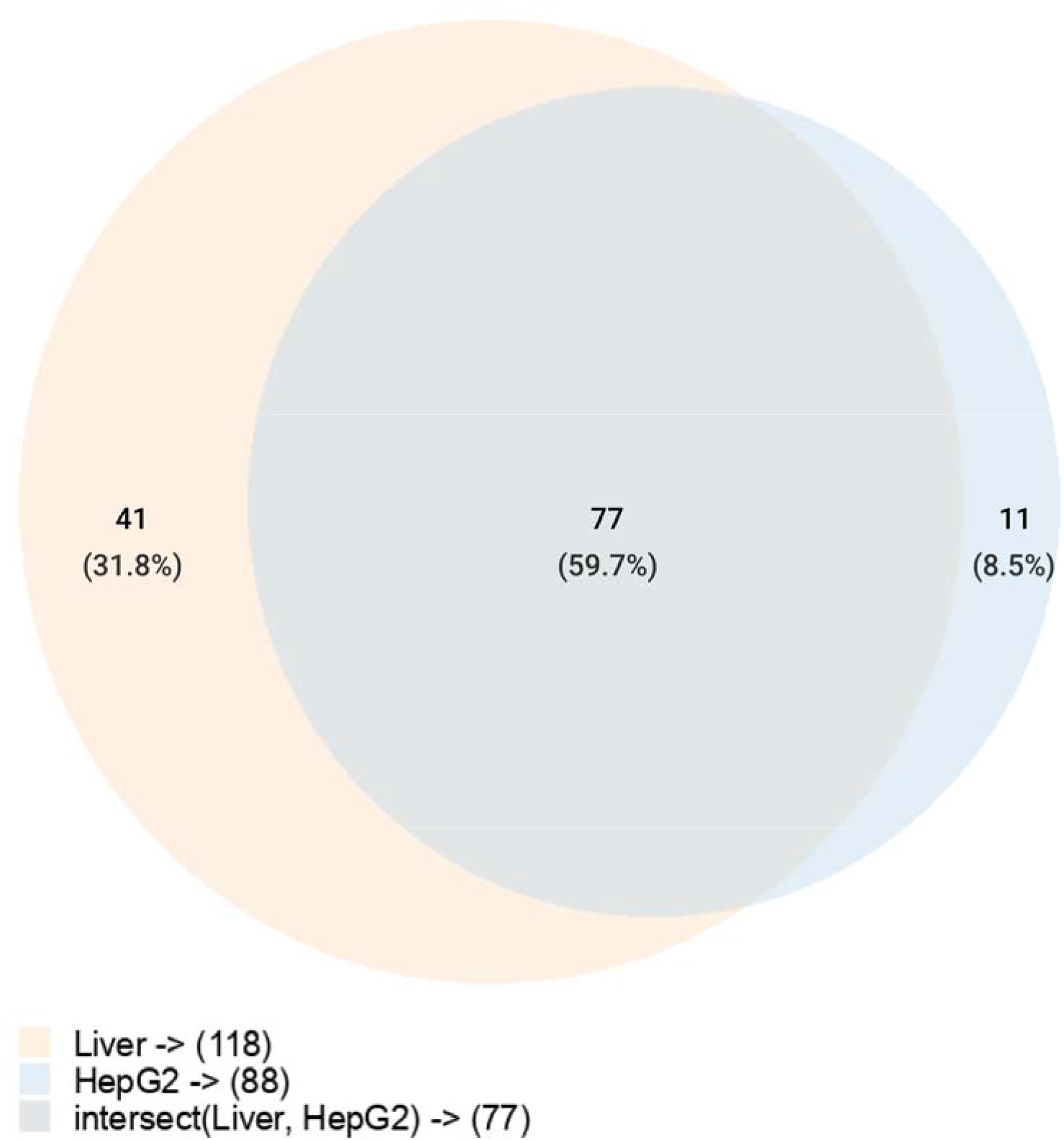
Comparison of chromosome 18 proteins identified in human liver cells and HepG2 cells using SRM analysis.

## 4. Materials and Methods

### 4.1 Liver samples collection

Human liver samples were collected at autopsy from three male donors, №1, №3, and №5, aged 65, 38, and 54 years old, respectively. We obtained the approval of the N.I. Pirogov Russian State Medical University Ethics Committee (protocol №3; March 15, 2018) and informed consent from the donor’s representatives. The donors were HIV- and hepatitis free, and the sections had no histological signs of liver disease. The postmortem resected samples were immediately placed in an RNAlater stabilization solution (Thermo Fisher Scientific, USA) and stored at −20 °C until further use [15].

### 4.2 Sample preparation

Liver tissue (0.1 mg) was lysed in a buffer containing 2 M urea, 1% deoxycholic acid, 15% acetonitrile, 147 mM NaCl, 10 mM TCEP, and 10 mM Na/Pi buffer (pH 7.4). The crude homogenate was sonicated, and the resulting solution was heated to 60 °C for 20 min. After cooling, the proteins were alkylated with 50 mM 2-CAA. The samples were diluted with 100 mM TEA (pH 8.0) to a total protein concentration of 1 mg/mL. Trypsin was added at a 1:100 w/w ratio, and proteins were digested at 38 °C overnight. The digestion was stopped by formic acid at a final concentration of 3%. Samples were centrifuged at 10000×g at 4 °C for 10 min. The peptide samples were evaporated in a vacuum concentrator and reconstituted in 0.5% formic acid.

Separation of peptides by RP chromatography under alkaline conditions and peptide fractions collection

Chromatographic separation of 200 μg of peptides under alkaline conditions (pH 9.0) and fraction collection of digested liver proteins was performed as described previously3. The collected fractions (750 μL) were dried in a vacuum at 30 °C. Dried fractions were reconstituted in 35 μL of 0.5% formic acid for subsequent analysis using LC-SRM and LC-MS/MS approaches.

### 4.3 Shotgun analysis

MS analysis was performed at least in triplicate using a Q Exactive HF-X mass spectrometer (Q Exactive HF-X Hybrid Quadrupole-Orbitrap mass spectrometer, Thermo Fisher Scientific, Rockwell, IL, USA) with the installed ESI-NSI ion source. The instrument was operated in positive ionization mode with emitter voltage adjusted to 2.0 kV and drying gas temperature at 280 °C. Surveyed in a range of 350 m/z–1500 m/z precursor ions (maximum injection time was 100 ms) with charge states from z = 2+ to z = 6+ were isolated in the quadrupole mass analyzer within ±2 m/z, and triggered to fragmentation in the m/z range with a fixed lower mass (140 m/z) and a dynamic upper mass (depending on the charge state of the fragmented precursor) limited to 2100 m/z. The data acquisition mode was set as Full MS/ddMS2, Top15, MS1 resolution – 60000 FWHM, AGC target – 1E6, MS2 resolution – 15000 FWHM, AGC target – 5E4, dynamic exclusion – 20 seconds. Liquid chromatography separation was accomplished on an Ultimate 3000 RSLCnano (Thermo Scientific, Rockford, IL, USA). Samples were loaded directly onto Acclaim Pepmap® C18 (75 μm × 150 mm, 2 μm) (Thermo Scientific, USA) analytical column at a flow rate of 1 μL/min for 15 minutes in 5% acetonitrile, supplied by 0.1% formic acid and 0.01% trifluoroacetic acid. Analytical separation was carried out at a flow rate 0.3 μL/min in a gradient of mobile phase A (water with 0.1% formic acid and mobile phase B (80% acetonitrile with 0.1% formic acid) in the following gradient: 2%–37% of mobile phase B for 60 min, followed by column washing in 90% of mobile phase B for 8 min, with equilibration of the column under initial gradient conditions (2% of mobile phase B) for 15 min before starting the next run. Data was processed using MaxQuant software v1.6.3.4 by target-decoy method [16]. The maximum number of missed cleavages allowed by trypsin-2. The oxidation of M and acetylation of protein N-term were set as variable modifications. Carbamidomethylation was set as a fixed modification. The latest release of the neXtProt database was used for protein identification (release 2021-02-15). Peptide spectrum matches (PSMs), peptides, and proteins were validated at a 1.0% false discovery rate (FDR) estimated using the decoy hit distribution; the FDRs of PSM, peptide, and protein were 0.12%, 0.63%, and 0.91%, respectively.

### 4.4 Internal standard synthesis

The SRM Atlas database was used to select the proteotypic peptides [17]. The peptides of choice were obtained using solid-phase peptide synthesis on the Overture (Protein Technologies, USA) or Hamilton Microlab STAR devices according to a published method [18]. Isotope-labeled leucine (Fmoc-Leu-OH 13C6, 15N), arginine (13C6, 15N4), lysine (13C6, 15N2), or serine (13C3, 15N1) were used for isotope-labeled peptide synthesis. The concentrations of the synthesized peptides were measured via amino acid analysis with fluorescent signal detection of amino acids following acidic hydrolysis of the peptides.

### 4.5 SRM SIS analysis

The selection of unique proteotypic peptides and the most intense transitions were performed based on the SRM scouting of chromosome 18 results. Reconstituted peptides were separated using an Infinity 1290 UPLC system (Agilent, Palo Alto, CA, USA), and peptides were detected using a triple-quadrupole G6495 mass spectrometer (Agilent, Palo Alto, CA, USA) in dynamic SRM SIS mode. The m/z of precursors, m/z of transition ions, CE values, b, y ions, transition ions, and retention time of the peptides on SIS and target peptides are presented in Table S1 and Table S2 (chromosome 18 transitions and UPS1 transitions, respectively). Each SRM experiment was triplicated. In each SRM experiment, 20 μg of peptides were used. Data obtained by the SRM SIS analysis were processed using the Skyline (version 20.1) software [19]. The results were manually inspected using the Skyline software to find transitions similar to those in the target peptides. Briefly, the peptide was considered to be detected in the run if the differences between relative intensities for three transitions of endogenous and isotopically labeled peptides did not exceed 25%, and the transition chromatographic profiles of endogenous peptides were identical to the corresponding transitions of stable isotope-labeled peptides. For each peptide, a calibration curve was built and the limit of detection (LOD) was determined in a pure standard solution. When analyzing the data, only those signals of endogenous peptides were accepted that exceeded the level of the minimum signal determined during calibration. If the signal was below the LOD but was reproduced in each repetition, such a signal was received. Quantitative analysis was based on the ratio of endogenous peptide peak area and stable isotope-labeled peptide peak area calculated using the Skyline software. Stable isotope-labeled peptides with known concentrations were added to the samples.

## 5. Conclusions

### Lessons learned from UPS1 analysis

The main problem with the proteomic analysis is the sensitivity of the instrument in the study of complex biological samples. Using a standard UPS1 set as an example, we revealed that the recovery of the sample from a large volume by any appropriate method at detectable concentrations solves the sensitivity problem. However, this technique cannot be applied when analyzing complex biological samples. In a biological sample of whole cell lysate, we could not select and study only the proteins encoded by chromosome 18 but should deal with the entire proteome. The analysis of UPS1 proteins revealed that to register all the proteins encoded by human chromosome 18, the presence of all proteins in the mixture must be at a concentration of 10^−9^ M or 1200 copies/cell.

Some positive effects can be achieved by fractionation under alkaline conditions. However, due to technical limitations (the maximum amount of protein that can be loaded on the Agilent Zorbax Poroshell 120 SB-C18 2.7 μm, 2.1 x 150 mm HPLC column), the amount of protein for separation can only be increased by 10 folds. Unlike UPS1, the concentration of proteins in a cell cannot be predicted in advance. The results showed that the loss of proteins begins at a concentration of 10^−10^ M, despite their presence in the sample. However, there is a certain benchmark that we can rely on in a biological sample, that is mRNA. The presence of mRNA molecules in a sample indicates the possible presence of the corresponding protein in the sample. However, data on the low correlation between mRNA and protein contents suggest that we cannot rely on 100% of the transcriptome [20, 21]. Some progress could be made in this direction within the framework of the study of the translatome, that is, only those RNA molecules that are bound to ribosomes and are most likely translated [22]. The convergence of proteomic and transcriptomic data means the detection of transcripts and their corresponding protein products. Our previous experience shows that the results of transcriptome analysis do not correlate with proteomic ones. Moreover, when examining the transcriptome by various bioinformatic methods, it is possible to detect the complete transcriptome of chromosome 18th genome in the liver. This does not mean that all transcripts are translated into proteins. In the future, we plan to use translatomic data for comparison with the proteome.

The application of all four analysis technologies (1D shotgun, 1D SRM, 2D shotgun, and 2D SRM) allowed the identification of 50% of the proteins encoded by Chr18 and achieving 50% coverage of the proteome registered. However, this level of sensitivity of proteomic technologies is not sufficient for identifying missing proteins in liver cells.

## Supplementary Materials

Table S1: Transitions for SRM analysis of proteins encoded by 18 human chromosome

Table S2: Transitions for SRM analysis of UPS1 proteins

Table S3: List of proteins identified by shotgun MS

Table S4: Calculated concentrations of proteins identified by 1D SRM (Chr 18)

Table S5: Calculated concentrations of proteins identified by 2D SRM,Figure S1: Venn diagram number of identifications by 1D and 2D shotgun in liver samples

Figure S2: Venn diagram showing intraindividual variability between liver samples

Figure S3: Diagram demonstrating gene coverage by each chromosome.

## Author Contributions

Conceptualization, Victor Zgoda and Alexander Archakov; Data curation, Olga Tikhonova; Formal analysis, Nikita Vavilov; Investigation, Nikita Vavilov and Olga Tikhonova; Methodology, Nikita Vavilov; Project administration, Alexander Archakov; Resources, Olga Tikhonova; Software, Ekaterina Ilgisonis and Andrey Lisitsa; Supervision, Andrey Lisitsa and Victor Zgoda; Visualization, Nikita Vavilov; Writing – original draft, Nikita Vavilov and Ekaterina Ilgisonis; Writing – review & editing, Andrey Lisitsa, Victor Zgoda and Alexander Archakov.All authors have read and agreed to the published version of the manuscript.

## Institutional Review Board Statement

The approval of the N.I. Pirogov Russian State Medical University Ethics Committee was obtained (protocol №3; March 15, 2018) and informed consent from the donor’s representatives.

## Data Availability Statement

Mass spectrometry data is available via ProteomeXchange with identifier PXD026997.

## Acknowledgments

This research was funded by RUSSIAN SCIENCE FOUNDATION, RSF, grant number 20-15-00410. (http://www.rscf.ru). The authors would like to thank the “Human Proteome” Core Facility, Institute of Biomedical Chemistry (IBMC), for the experimental work performed.

